# Crop domestication and pathogen virulence: Interactions of tomato and *Botrytis* genetic diversity

**DOI:** 10.1101/255992

**Authors:** Nicole E. Soltis, Susanna Atwell, Gongjun Shi, Rachel Fordyce, Raoni Gwinner, Dihan Gao, Aysha Shafi, Daniel J. Kliebenstein

## Abstract

Human selection during crop domestication alters numerous traits, including disease resistance. Studies of qualitative resistance to specialist pathogens typically find decreased resistance in domesticated crops in comparison to their wild relatives. However, less is known about how crop domestication affects quantitative interactions with generalist pathogens. To study how crop domestication impacts plant resistance to generalist pathogens, and correspondingly how this interacts with the pathogen’s genetics, we infected a collection of wild and domesticated tomato accessions with a genetically diverse population of the generalist pathogen *Botrytis cinerea*. We quantified variation in lesion size of 97 *B. cinerea* genotypes (isolates) on 6 domesticated *Solanum lycopersicum* and 6 wild *S. pimpinellifolium* genotypes. This showed that lesion size was significantly controlled by plant domestication, plant genetic variation, and the pathogen’s genotype. Overall, resistance was slightly elevated in the wild germplasm in comparison to domesticated tomato accessions. Genome-wide association (GWA) mapping in *B. cinerea* identified a highly polygenic collection of genes. This suggests that breeding against this pathogen would need to utilize a diversity of isolates to capture all possible mechanisms. Critically, we identified a discrete subset of *B. cinerea* genes where the allelic variation was linked to altered virulence against the wild versus domesticated tomato accessions. This indicates that this generalist pathogen already has the necessary allelic variation in place to handle the introgression of wild resistance mechanisms into the domesticated crop. Future studies are needed to assess how these observations extend to other domesticated crops and other generalist pathogens.

## Introduction

Plant disease is mediated by complex interactions among diverse host and pathogen molecular pathways, and the disease outcome is the sum of host plant susceptibility/resistance and pathogen virulence/sensitivity mechanisms. The specific outcome of any interaction is highly dependent on the genetic variation within these pathways in both the host and pathogen. Over time, mutation and selection have led to distinct genetic architectures in the host and pathogen that are at least partly influenced by the host range of the pathogen. Specialist pathogens are a major focus in plant pathology; virulent on a narrow range of hosts, and often limited to a single species or genus. Most known plant genes for resistance to specialist pathogens confer qualitative resistance through innate immunity via large-effect loci that enable the recognition of the pathogen (Dangl and Jones 2001, Jones and Dangl 2006, Dodds and Rathjen 2010, Pieterse, Van der Does et al. 2012). These recognition signals can be conserved pathogen patterns such as cell-wall polymers or flagellin, or alternatively, specific virulence factors that block perception of the pathogen, and in turn are detected by plant proteins that guard the signaling networks (Jones and Dangl 2006, Bittel and Robatzek 2007, Ferrari, Galletti et al. 2007, Boller and He 2009, Dodds and Rathjen 2010). The evolution of large-effect qualitative loci has partly been driven by the narrow host range for the pathogen that enhances co-evolution between host resistance genes and pathogen virulence mechanisms.

In contrast to specialist pathogens, generalist pathogens are virulent across a wide range of plant host species. Generalist pathogens potentially have less stringent co-evolution to specific hosts and their accompanying resistance mechanisms, because these pathogens can easily shift to new hosts in the environment. This allows generalist pathogens to evade the rapid evolution of new resistance mechanisms within specific hosts until they evolve to counter this new resistance. This niche-shifting ability may partially explain the observation that most natural resistance to generalist pathogens is highly polygenic, and the underlying plant genes for resistance are quantitative (Glazebrook 2005, Nomura, Melotto et al. 2005, Goss and Bergelson 2006, Rowe and Kliebenstein 2008, Barrett, Kniskern et al. 2009, Corwin, Copeland et al. 2016). Plant quantitative resistance genes to generalist pathogens include a broad array of direct defense genes, like those involved in secondary metabolite production, cell wall formation, and defense proteins (Zhang, Khan et al. 2002, Denby, Kumar et al. 2004, Zipfel, Robatzek et al. 2004, Ferrari, Galletti et al. 2007, Rowe and Kliebenstein 2008, Poland, Balint-Kurti et al. 2009, Corwin, Copeland et al. 2016). Importantly, these quantitative plant resistance loci do not alter resistance to all genotypes (isolates) of a pathogen but interact with the infecting pathogen’s genotype. For example, the ability of the *Arabidopsis* defense metabolite, camalexin, to provide resistance to *Botrytis cinerea* depends upon whether the specific isolate is sensitive or resistant to camalexin (Kliebenstein, Rowe et al. 2005, Pedras and Ahiahonu 2005, Stefanato, Abou‐Mansour et al. 2009, Pedras, Hossain et al. 2011) and similarly *B. cinerea* virulence on tomato varies with the isolate’s ability to detoxify tomatine (Quidde, Osbourn et al. 1998, Quidde, Büttner et al. 1999). In contrast to the polygenic nature of plant resistance to generalist pathogens, little is known about the genetic architecture of virulence within generalist pathogens, and how this is affected by genetic variation in the plant (Bartoli and Roux 2017). There are no reported naturally variable large-effect virulence loci in generalist pathogens, suggesting that virulence in generalist pathogens is largely quantitative and polygenic. This potential for interaction between polygenic virulence in generalist pathogens and equally polygenic resistance in host plants suggests that we need to work with genetic variation in both the host and pathogen to truly understand quantitative host-pathogen interactions.

A key evolutionary process in plants that has affected resistance to specialist pathogens is the domestication of crop plants. Domesticated plant varieties are typically more sensitive to specialist pathogens than their wild relatives (Smale 1996, Rosenthal and Dirzo 1997, Couch, Fudal et al. 2005, Dwivedi, Upadhyaya et al. 2008), and pathogens may evolve higher virulence on domesticated hosts (Stukenbrock and McDonald 2008). Further, domestication typically imposes a genetic bottleneck that reduces genetic diversity in the crop germplasm, including decreased availability of resistance alleles against specialist pathogens (Tanksley and McCouch 1997, Doebley, Gaut et al. 2006, Chaudhary 2013). These general evolutionary patterns, of lower resistance and allelic diversity found when studying the interaction of specialist pathogens with crop plants, are assumed to similarly hold for generalist pathogens and their domesticated hosts. However, there is less information about how crop host domestication affects disease caused by generalist pathogens, when the resistance to these pathogens is quantitative and polygenic rather than qualitative and monogenic. As such, there is a need to conduct a detailed analysis of how domestication may alter the interaction of a plant with a broad generalist pathogen, and correspondingly, how domestication influences the pathogen.

*Botrytis cinerea* provides a model generalist pathogen for studying quantitative interactions with plant hosts, and underlying evolutionary processes for this generalist in contrast to specialist pathogens. *B. cinerea* is a broad generalist pathogen that can infect most tested plants from bryophytes to eudicots, and causes wide ranging pre‐ and post-harvest crop losses (Nicot and Baille 1996, Elad, Williamson et al. 2007, Fillinger and Elad 2015). Individual isolates of *B. cinerea* show the same broad host range (Deighton, Muckenschnabel et al. 2001, Finkers, van Heusden et al. 2007, Ten Have, van Berloo et al. 2007, Corwin, Subedy et al. 2016), in contrast to pathogens like *Fusarium oxysporum* where the species can infect diverse hosts, but each isolate is highly host specific (Katan 1999, Ormond, Thomas et al. 2010, Loxdale, Lushai et al. 2011, Barrett and Heil 2012). *B. cinerea* isolates display significant variation in virulence phenotypes, partly due to genetic variation in specific virulence mechanisms, like the production of the phytotoxins, botrydial and botcinic acid (Siewers, Viaud et al. 2005, Dalmais, Schumacher et al. 2011). This genetic variation also influences cell wall degrading enzymes and key regulators of virulence like *VELVET* that quantitatively control virulence on multiple host plants (Rowe and Kliebenstein 2007, Schumacher, Pradier et al. 2012). This genetic variation in diverse virulence mechanisms can contribute to the formation of quantitative differences in virulence between the isolates (ten Have, Mulder et al. 1998). The phenotypic variation is driven by a high level of sequence diversity spread across the genome (Rowe and Kliebenstein 2007, Fekete, Fekete et al. 2012, Atwell, Corwin et al. 2015, Atwell, Soltis et al. 2017). The polymorphism rate in *B. cinerea* was measured as 6.6 SNP/kb, which is more variable than most previously studied plant pathogens (1-2 SNP/kb in *Blumeria graminis*, 1.5 SNP/kb in *Melampsora larici-populina*, 5.5 SNP/kb in the compact genome of the obligate biotroph *Plasmodiophora brassicae*), and close to the genetic diversity found in the human pathogen *Mycobacterium tuberculosis* (2.9 to 6.2 SNP/kb) (Farhat, Shapiro et al. 2013, Hacquard, Kracher et al. 2013, Wicker, Oberhaensli et al. 2013, Persoons, Morin et al. 2014, Desjardins, Cohen et al. 2016, Power, Parkhill et al. 2017). Higher polymorphism rates are reported for the wheat stem rust pathogen *Puccinia graminis* f. sp. *tritici*, from a small non-random sample of isolates (12.3 SNP/kb) (Upadhyaya, Garnica et al. 2014). In addition to SNP diversity, the genomic sequencing showed that *B. cinerea* has a high level of recombination and genomic admixture, as if it were a randomly intermating population. As such, a collection of *B. cinerea* isolates contains genetic variation in a wide range of virulence mechanisms, offering the potential to challenge the host with a blend of diverse virulence mechanisms. This can potentially identify the pathogen variation controlling quantitative virulence, even in non-model plant systems (Bartoli and Roux 2017).

A model pathosystem for studying quantitative host-pathogen interactions during domestication is the tomato-*B. cinerea* system, where the pathogen causes crop loss due to both pre‐ and post-harvest infection (Dean, Van Kan et al. 2012, Hahn 2014, Romanazzi and Droby 2016). Resistance to *B. cinerea* is a quantitative trait in tomato as with most other species, with identified tomato QTLs each explaining up to 15% of phenotypic variation for lesion size on stems (Dıaz, ten Have et al. 2002, Finkers, van Heusden et al. 2007, Ten Have, van Berloo et al. 2007, Rowe and Kliebenstein 2008, Corwin, Copeland et al. 2016). Tomato is also a key model system to study how domestication influences plant physiology and resistance, including alterations in the circadian clock (Tanksley 2004, Bai and Lindhout 2007, Panthee and Chen 2010, Bergougnoux 2014, Müller, Wijnen et al. 2016), which can modulate resistance to *B. cinerea* (Sauerbrunn and Schlaich 2004, Weyman, Pan et al. 2006, Bhardwaj, Meier et al. 2011, Hevia, Canessa et al. 2015). This suggests that host plant domestication within tomato can alter traits known to influence *B. cinerea* resistance from other systems. Thus we are using the tomato-*B. cinerea* pathosystem to directly measure the interaction of crop domestication with genetic variation in a generalist pathogen to better understand the evolution of this pathosystem.

In this study, we infected 97 genetically diverse *B. cinerea* isolates on a collection of domesticated tomato, *S. lycopersicum*, and wild tomato, *S. pimpinellifolium*, and quantified the interaction through lesion size in a detached leaf assay. Previous studies have examined *B. cinerea* resistance between domesticated and distantly related wild tomato species (i.e. *S. lycopersicum* and *S. pimpinellifolium*) using single isolates of pathogens (Egashira, Kuwashima et al. 2000, Nicot, Moretti et al. 2002, Guimaraes, Chetelat et al. 2004, Ten Have, van Berloo et al. 2007, Finkers, Bai et al. 2008). These previous studies typically used individual wild and domesticated tomato accessions that were the founders of mapping populations, and found a wide range of *B. cinerea* resistance. However, it is still unknown how domesticated and closely related wild tomatoes compare for *B. cinerea* resistance using multiple plant genotypes and a population of the pathogen. In this study, we asked whether *B. cinerea* virulence was controlled by host variation, pathogen variation, or the interaction between them. Lesion size of *B. cinerea* is a quantitative trait that was controlled by plant domestication status, plant genotype and pathogen isolate. We looked for evidence of specialization within our generalist pathogen population. While our *B. cinerea* isolates appear to be generalists across domestication in *Solanum,* a subset of isolates are sensitive to tomato domestication. Finally, we aimed to identify the genetic basis of variation in *B. cinerea* virulence on domesticated and wild tomato. We conducted genome-wide association (GWA) in *B. cinerea* to identify pathogen loci where genetic variation leads to altered virulence across the host genotypes, including a specific test for loci that influence responses to crop domestication. Few studies have conducted GWA in plant pathogens for virulence phenotypes, and most of these were limited by few variable loci or few genetically distinct isolates (Dalman, Himmelstrand et al. 2013, Gao, Liu et al. 2016, Talas, Kalih et al. 2016, Wu, Sakthikumar et al. 2017). At the genetic level, virulence of *B. cinerea* is highly quantitative, with hundreds of significant SNPs with small effect sizes associated with lesion area on each tomato genotype. Importantly, there is a subset of loci in the pathogen where allelic variation gives the isolates opposing responses to crop domestication. These pathogen loci could provide tools for understanding how domestication in tomato has influenced generalist pathogen resistance, to inform breeding efforts.

## Methods

### Tomato genetic resources

We obtained seeds for 12 selected tomato genotypes in consultation with the UC Davis Tomato Genetics Resource Center. These include a diverse sample of 6 genotypes of domesticated tomato’s closest wild relative (*S. pimpinellifolium*) from throughout its native range (Peru, Ecuador) and 6 heritage and modern varieties of *S. lycopersicum*. We bulked all genotypes in long-day (16h photoperiod) greenhouse conditions at UC Davis in fall 2014. We grew plants under metal-halide lamps using day/night temperatures at 25°C/18°C in 4” pots filled with standard potting soil (Sunshine mix #1, Sun Gro Horticulture). Plants were watered once daily and pruned and staked to maintain upright growth. Fruits were collected at maturity and stored at 4°C in dry paper bags until seed cleaning. To clean the seeds, we incubated seeds and locule contents at 24°C in 1% protease solution (Rapidase C80 Max) for 2h, then rinsed them in deionized water and air-dried. We then stored seeds in a cool, dry, dark location until use.

To grow plants for detached leaf assays, we bleach-sterilized all seeds and germinated them on paper in the growth chamber using flats covered with humidity domes. At 7 days we transferred seedlings to soil (SunGro Horticulture, Agawam, MA) and grew all plants in growth chambers in 20°C, short-day (10h photoperiod) conditions with 180-190 uM light intensity and 60% RH. We bottom-watered with deionized water every two days for two weeks, and at week 3 watered every two days with added nutrient solution (0.5% N-P-K fertilizer in a 2-1-2 ratio; Grow More 4-18-38). The plants were used for detached leaf assays 6 weeks after transferring seedlings to soil.

### *B. cinerea* genetic resources

We utilized a previously described collection of *B. cinerea* isolates that were isolated as single spores from natural infections of fruit and vegetable tissues collected in California and internationally (Atwell, Corwin et al. 2015, Zhang, Corwin et al. 2017). This included five isolates obtained from natural infections of tomato. We maintained *B. cinerea* isolates as conidial suspensions in 30% glycerol for long-term storage at −80°C. For regrowth, we diluted spore solutions to 10% concentration in filter-sterilized 50% grape juice, and then inoculated onto 39g/L potato dextrose agar (PDA) media. We grew isolates at 25°C in 12h light, and propagated every 2 weeks. Sequencing failed for 6 out of our 97 phenotyped isolates. For GWA mapping with the 91 isolates genotyped in this study, we utilized a total of 272,672 SNPs with minor allele frequency (MAF) 0.20 or greater, and less than 10% missing calls across the isolates (SNP calls in at least 82/ 91 isolates).

### Detached leaf assay

To study the effect of genetic variation in host and pathogen on lesion formation, we infected detached leaves of 12 diverse tomato varieties with the above 97 *B. cinerea* isolates. We used a randomized complete block design for a total of 6 replicates across 2 experiments. In each experiment, this included a total of 10 plants per genotype randomized in 12 flats in 3 growth chambers. Each growth chamber block corresponded with a replicate of the detached leaf assay, such that growth chamber and replicate shared the same environmental block. At 6 weeks of age, we selected 5 leaves per plant (expanded leaves from second true leaf or older), and 2 leaflet pairs per leaf. We randomized the order of leaves from each plant, and the leaflets were placed on 1% phytoagar in planting flats, with humidity domes. Our inoculation protocol followed previously described methods (Denby, Kumar et al. 2004, Kliebenstein, Rowe et al. 2005). Spores were collected from mature *B. cinerea* cultures grown on canned peach plates, and diluted to 10 spores/ µL in filter-sterilized 50% organic grape juice. 4µl droplets of the diluted spore suspensions were placed onto the detached leaflets at room temperature. Mock-inoculated control leaves were treated with 4µL of 50% organic grape juice without spores. Digital photos were taken of all leaflets at 24, 48, and 72 hours post inoculation and automated image analysis was used to measure lesion size.

### Automated Image Analysis

Lesion area was digitally measured using the EBImage and CRImage packages (Pau, Fuchs et al. 2010, Failmezger, Yuan et al. 2012) in the R statistical environment (R Development Core Team 2008), as previously described (Corwin, Copeland et al. 2016, Corwin, Subedy et al. 2016). Leaflets were identified as objects with green hue, and lesions were identified as low-saturation objects within leaves. Images masks were generated for both the leaf and lesion, then manually refined by a technician to ensure accurate object calling. The area of these leaves and lesions were then automatically measured as pixels per lesion and converted to area using a 1 cm reference within each image.

### Data analysis

We analyzed lesion areas using a general linear model for the full experiment, including the fixed effects of isolate genotype, plant domestication (*S. lycopersicum* or *S. pimpinellifolium*), plant genotype (which is nested within domestication status), experiment, and block (nested within experiment) on lesion area, as well as their interactions (lme4; (Douglas Bates 2015)). Two of our 97 isolates that did not have replication across 2 experiments were dropped at this stage of analysis. There was no difference in the results if experiment and block were treated as random effects. Adding terms for individual plant, leaf, and leaflet position did not significantly improve the full model, so they were omitted from further analysis. We also tested a mixed model with random effects of experiment and block, but this did not affect our interpretation of the fixed effects. This model was used to calculate the significance of each factor and to obtain the least-squared means of lesion size for each *B. cinerea* isolate x tomato accession as well as for each *B. cinerea* isolate x domestic/wild tomato. We also calculated a domestication sensitivity phenotype, Sensitivity = (Domesticated lesion size – Wild lesion size) / Domesticated lesion size.

We used several methods to examine host specialization to tomato within *B. cinerea*. First, we split our *B. cinerea* population into isolates collected from tomato tissue vs. other hosts. We compared these groups by t-test for virulence on domesticated tomato genotypes, wild tomato genotypes, or all tomato genotypes. Next, we used a Wilcoxon signed-rank test to compare the rank order distribution of lesion sizes across paired tomato genotypes. To examine host specialization to tomato domestication within *B. cinerea*, we used a Wilcoxon signed-rank test to compare the rank order of lesion sizes across all domesticated vs. all wild tomato genotypes. Finally, we conducted single-isolate ANOVAs with FDR correction to identify isolates with a significant response to plant genotype or domestication status.

The model means and Sensitivity were used as the phenotypic input for GWA using bigRR, a heteroskedastic ridge regression method that incorporates SNP-specific shrinkage (Shen, Alam et al. 2013). This approach has previously had a high validation rate (Ober, Huang et al. 2015, Corwin, Copeland et al. 2016, Francisco, Joseph et al. 2016, Kooke, Kruijer et al. 2016). The *B. cinerea* GWA used 272,672 SNPs at MAF 0.20 or greater and <10% missing SNP calls as described above. Because bigRR provides an estimated effect size, but not a p-value, significance was estimated using 1000 permutations to determine effect significance at 95%, 99%, and 99.9% thresholds (Doerge and Churchill 1996, Shen, Alam et al. 2013, Corwin, Copeland et al. 2016). SNPs were annotated using SNPdat (Doran and Creevey 2013) with gene transfer format file construction from the T4 gene models for genomic DNA by linking the SNP to genes within a 2kbp window (<u>http://www.broadinstitute.org</u>, (Staats and van Kan 2012)). Functional annotations are based on the T4 gene models for genomic DNA (http://www.broadinstitute.org, *B. cinerea*; (Staats and van Kan 2012)). Additional genes of interest, based on a broad literature search of known virulence loci, were taken from NCBI (https://www.ncbi.nlm.nih.gov/) and included by mapping sequence to the T4 reference using MUMmer v3.0 (Kurtz, Phillippy et al. 2004). We used the program InterProScan within BLAST2GO for functional gene ontology (GO) annotation of the gene models (http://www.blast2go.com).

To predict expected overlap of significant SNPs across plant genotypes, we used the average number of significant SNPs per each of the 12 plant genotypes (14,000 SNPs) and calculated expected overlap between those 12 lists using binomial coefficients.

### Reagent and Data Availability

Supplemental information includes table S1 – S3. Table S1 shows mean lesion size of each isolate on each tomato plant genotype. Table S2 shows isolate rank order shifts across each pair of host plant genotypes. Table S3 includes gene annotation information for the top SNPs.

## Results

### Experimental Design

To measure how tomato domestication affects quantitative resistance to a population of a generalist pathogen, we infected a collection of 97 diverse *B. cinerea* isolates (genotypes) on wild and domesticated tomato genotypes. We compared domesticated and closely related wild tomatoes for *B. cinerea* resistance using multiple plant genotypes and a population of the pathogen. We selected 6 domesticated *Solanum lycopersicum* and 6 wild *S. pimpinellifolium* accessions, the closest wild relative of *S. lycopersicum*, to directly study how domestication has influenced resistance to *B. cinerea* (Peralta, Spooner et al. 2008, Müller, Wijnen et al. 2016). Our previously collected *B. cinerea* sample includes 97 isolates obtained from various eudicot plant hosts, including tomato stem tissue (2 isolates; T3, KT) and tomato fruit (3 isolates; KGB1, KGB2, Supersteak)(Atwell, Soltis et al. 2017). We infected all 97 *B. cinerea* isolates onto each of the 12 plant genotypes in 3-fold replication across 2 independent experiments in a randomized complete block design, giving 6 measurements per plant-pathogen combination, for a total of 3,276 lesions. Digital measurement of the area of the developing lesion provides a composite phenotype controlled by the interaction of host and pathogen genetics. This measurement of the plant-*B. cinerea* interaction has been used successfully in a number of molecular and quantitative genetic studies (Ferrari, Plotnikova et al. 2003, Denby, Kumar et al. 2004, Kliebenstein, Rowe et al. 2005, Ferrari, Galletti et al. 2007, Ten Have, van Berloo et al. 2007, AbuQamar, Chai et al. 2008, Rowe and Kliebenstein 2008, Liu, Hong et al. 2014).

### Lesion size (phenotypic) variation

We collected images of all lesions at 24, 48, and 72 hours post inoculation. At 24 hours, no visible lesions were present on the tomato leaves. At 48 hours, a thin ring of primary lesion becomes visible surrounding the location of the spore droplet, but no expansion is visible. At 72 hours significant lesion growth was visible, but no lesions had spread to infect over half of the leaflet. We digitally measured the area of all developing lesions at 72 hours post infection (HPI) as a measure of virulence (Figure 1). We observed a mean lesion size of 0.67 cm^2^ across the full experiment, with 0.94 CV across the full isolate population on all tomato genotypes. Individual isolates were highly variable in their lesion size across tomato genotypes (Figure 1 c-h), with mean lesion size per isolate of 0.14 cm^2^ to 1.29 cm^2^, and individual isolate coefficient of variation CV from 0.51 to 1.68 across all observations on all tomato genotypes (Table S1). A subset of these isolates are highly virulent on tomato (mean lesion size > 1.05 cm^2^, Figure 1e), and a subset can be considered saprophytic (mean lesion size < 0.3 cm^2^, Figure 1f).

**Figure 1.**
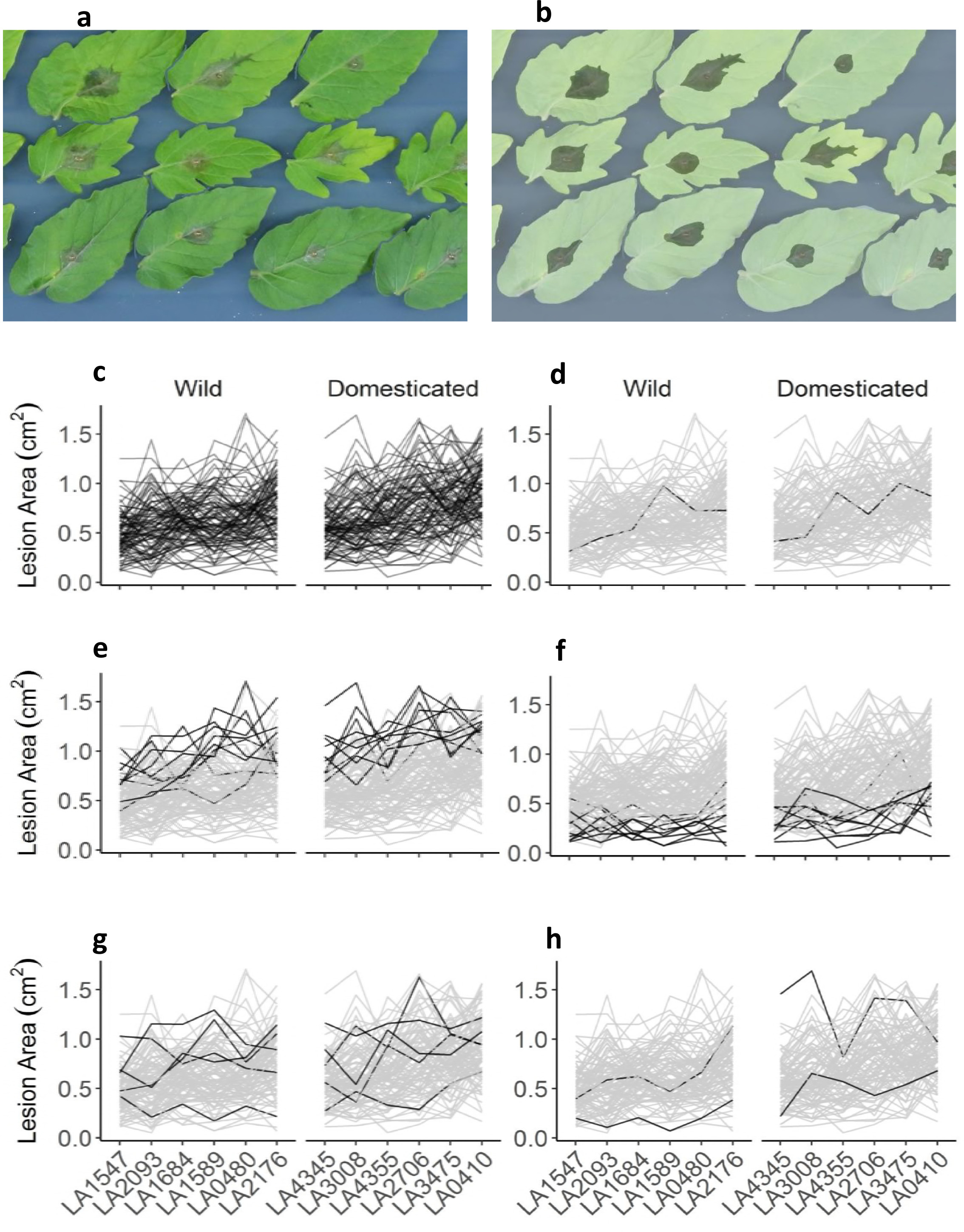
*Botrytis cinerea* x tomato diversity in detached leaf assay and digital image analysis. a) Individual tomato leaflets of 6 *S. lycopersicum* genotypes and 6 *S. pimpinellifolium* genotypes are in randomized rows, spore droplets of individual *B. cinerea* isolates are in randomized columns. Digital images are collected 72 hours post inoculation. Single droplets of 40 *B. cinerea* spores are infected on randomized leaflets using randomized isolates, and digital images are taken 72 hours post inoculation. b) Digital masking of leaf and lesion is followed by automated measurement of area for each lesion. c) Shown is an interaction plot of lesion size due to all individual *B. cinerea* isolates on all of the tomato host genotypes, grouped by domestication status. The x-axis includes each tomato host genotype. Each line traces the average lesion size of a single *B. cinerea* isolate across hosts. d) The common reference *B. cinerea* isolate B05.10 is highlighted in black. e) The ten highest-virulence isolates, as estimated by mean virulence across all tomato genotypes, are highlighted in black. f) The ten most saprophytic, or low virulence, isolates, as estimated by mean virulence across all genotypes, are highlighted in black. g) The five isolates collected from tomato tissue are highlighted in black. h) The two isolates with significant domestication sensitivity are shown in black.

### Contribution of Pathogen Genetics, Plant Genetics and Crop Domestication Effects on Resistance

To measure the relative contribution of genetic diversity in the plant and the pathogen to variation in the virulence/ susceptibility phenotype, we used a multiple linear regression model (R Development Core Team 2008). This model directly tested the contribution of plant genotype, plant domestication status, and pathogen genotype (isolate) to variation in lesion size. The final model explained 60% of the total variance for lesion size, and showed that genetic variation within both the host plant and the pathogen had significant effects on lesion growth, with pathogen isolate diversity explaining 3.5 fold more variance than plant genotype, 46% of total genetic variance for pathogen isolate vs. 13% for plant genotype (Table 1 and Figure 1c). Interestingly, tomato domestication status significantly impacted *B. cinerea* virulence, as shown by the small but significant effects of genetic variation between domesticated and wild tomatoes (3.5% of total genetic variance, Table 1). There was no evidence for significant interaction effects between pathogen isolate and plant genotype, but this term contributed the largest proportion of the plant-related variance in lesion size (34% of total genetic variance, Table 1). The lack of significance for this term in face of the large fraction of variance may be due to the vast degrees of freedom in this term (Table 1). Thus, the interaction between tomato and *B. cinerea* was significantly controlled by genetic diversity within the host plant and the pathogen, including a slight effect of domestication status.

**Table 1.**
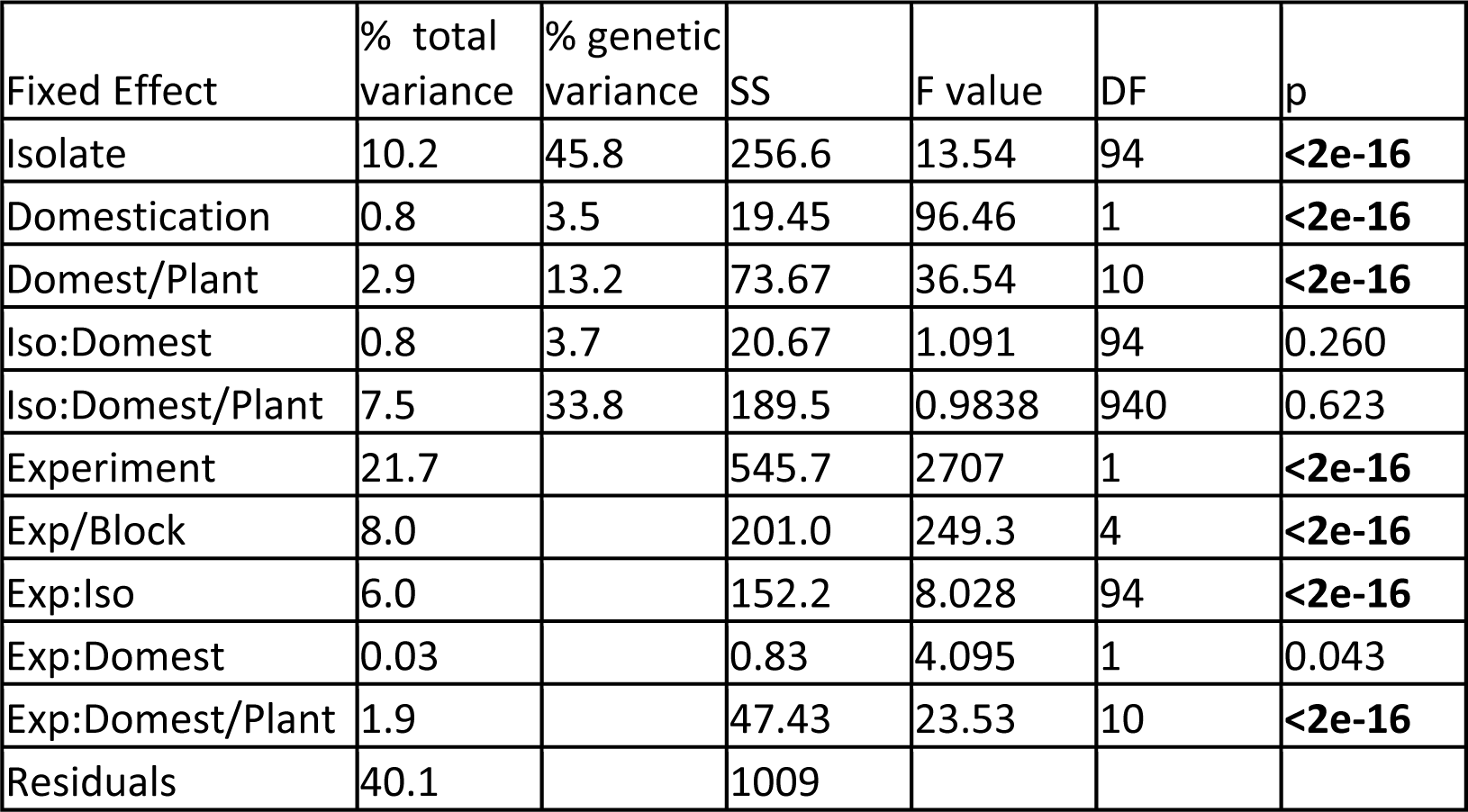
ANOVA results of the interaction between 12 tomato accessions and 95 *B. cinerea* isolates measured as lesion area. The Type III Sums-of-Squares, F-value, Degrees of Freedom and p-value for the linear modelling of lesion area for 12 tomato accessions by 95 *B. cinerea* isolates is shown. Two of our 97 isolates did not have replication across 2 experiments, so they were dropped at this stage of analysis. The terms are as follows; Isolate is the 95 *B. cinerea* isolates, Domestication is wild tomato, *S. pimpinellifolium*, versus domestic tomato, *S. lycopersicum*, Plant is 12 tomato genotypes nested within their respective domestication groupings, Experiment tests the 2 independent replicate experiments, Experiment/Block tests the three blocks nested within each experiment. In addition interactions of these factors are tested (:).

### Domestication and Lesion Size Variation

Existing literature predominantly reports that crop domestication decreases plant resistance to pathogens (Smale 1996, Rosenthal and Dirzo 1997, Couch, Fudal et al. 2005, Dwivedi, Upadhyaya et al. 2008, Stukenbrock and McDonald 2008). In our analysis, we identified a significant increase, 18%, in the resistance of wild tomato in comparison to domesticated tomato across the population of *B. cinerea* isolates (Figure 2, Table 1). However, this domestication effect was not the dominant source of variation, as genetic variation within the domesticated and wild genotypes contributed 3.8 fold more variation in resistance than domestication alone (Table 1). So while we did observe the expected decreased resistance in domesticated tomato, domestication was a minor player in controlling lesion size variation, with most of the plant genetic signature coming from variation within both the wild and domesticated tomato species.

**Figure 2.**
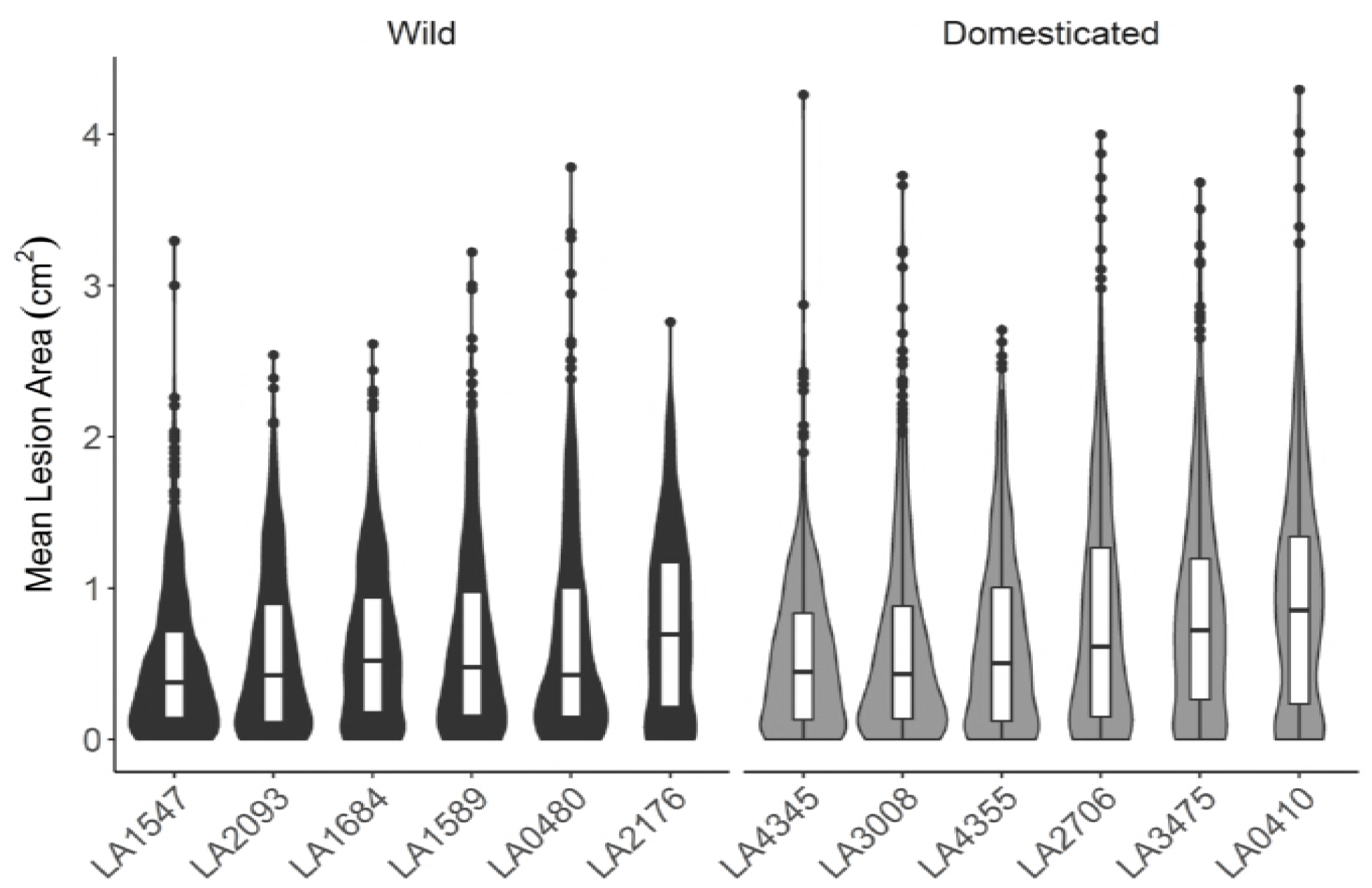
Distribution of tomato genotype susceptibility to infection with 97 genetically diverse *B. cinerea* isolates. Violin plots show the distribution of lesion size caused by *B. cinerea* isolates on each tomato host genotype. Individual points are mean lesion size for each of the 97 different isolate-host pairs. The boxes show the 75^th^ percentile distribution, and the horizontal line shows the mean resistance of the specific host genotype. The tomato genotypes are grouped based on their status as wild or domesticated germplasm.

In addition to altering trait means, domestication commonly decreases genetic variation in comparison to wild germplasm due to bottlenecks, including for tomato (Tanksley and McCouch 1997, Doebley, Gaut et al. 2006, Bai and Lindhout 2007). This decreased genetic variation should also limit phenotypic variation, including disease phenotypes. Interestingly in this tomato population, we did not observe reduced variation in lesion size in the wild tomato. Indeed, the domesticated tomato genotypes had a wider range of average lesion size than wild genotypes; the 90^th^ percentile range (95^th^ percentile to 5^th^ percentile) was 2.03 cm^2^ lesion size variation on domesticated tomato (standard deviation = 0.68 cm^2^) versus 1.76 cm^2^ variation on wild tomato (standard deviation = 0.58 cm^2^). Additionally, the wild and domesticated tomato genotypes showed statistically similar variation in resistance (F-test, F_96,96_=1.39, p=0.11)(Figure 3). Overall, there is a slight domestication impact on average resistance to *B. cinerea*, but no evidence of a phenotypic bottleneck due to domestication.

**Figure 3.**
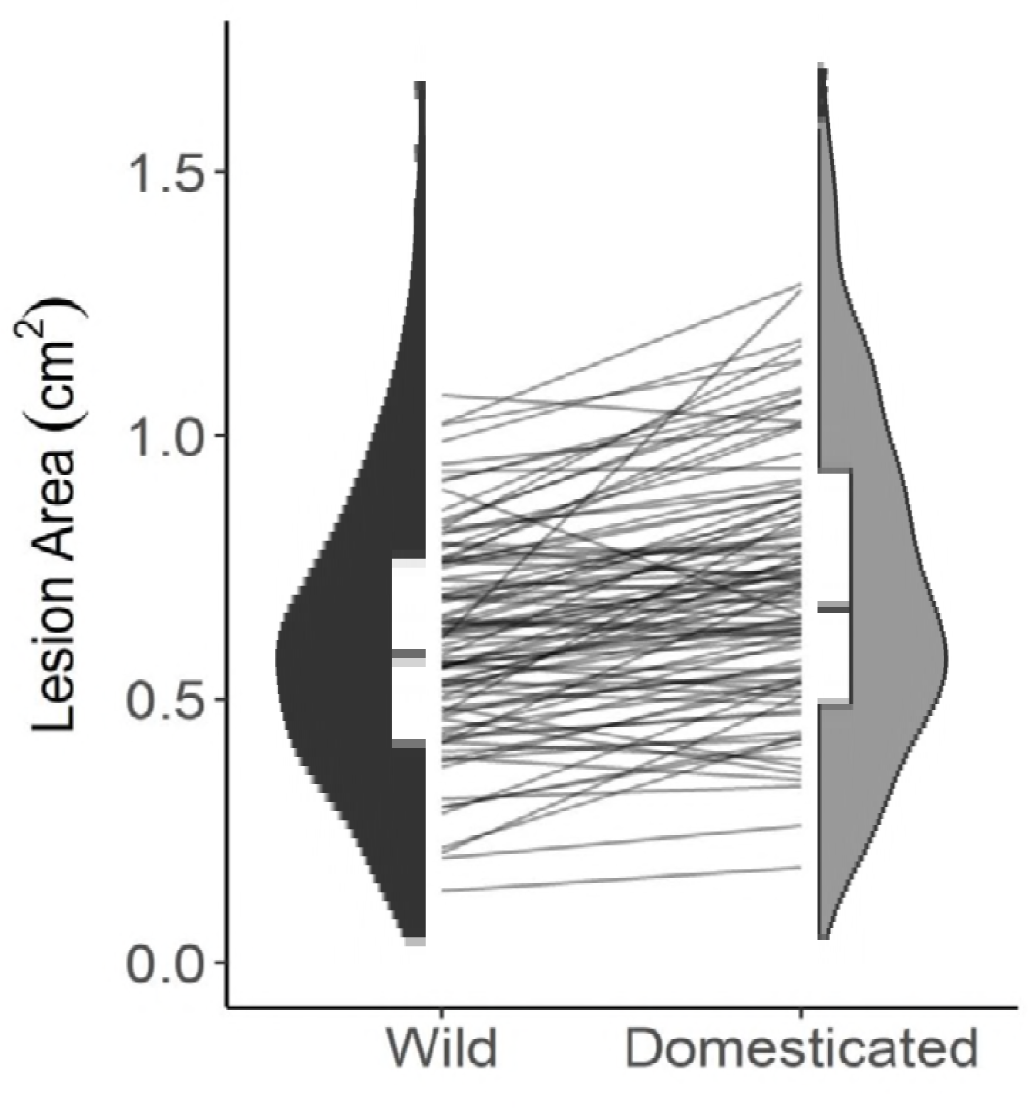
Distribution of *B. cinerea* virulence by tomato domestication status. The violin plots show the mean virulence of each *B. cinerea* isolate on the tomato genotypes, grouped as wild or domesticated germplasm. The domestication effect on lesion size is significant (Table 1 ANOVA, p<2e-16). The interaction plot between the two violin plots connects the average lesion size of a single *B. cinerea* isolate between the wild and domesticated germplasm.

### Pathogen Specialization to Source Host

One evolutionary model of generalist pathogens suggests that isolates within generalist pathogen species may specialize on specific hosts. Alternatively, isolates may also be generalists, with specialization absent even between individuals. Our collection of *B. cinerea* includes five isolates which may be adapted to tomato, as they were collected from *S. lycopersicum*. To test if there is evidence for specialization to the source host, we compared the virulence of the *B. cinerea* isolates obtained from tomato to the broader pathogen population. For *B. cinerea* genotypes isolated from tomato tissue vs. other hosts, there was no significant difference in lesion size on domesticated tomato (t-test; t=1.10, n = 97, p=0.33), wild tomato (t-test; t=1.09, n = 97, p=0.33) or across all tomato genotypes (t-test; n = 97, p=0.14) (Figure 1g). In fact, one isolate collected from tomato tissue (KGB1) was within the 10 least-virulent isolates and another (Triple3) was within the 10 most-virulent isolates (Figure 1g). This demonstrated significant genetic variation in virulence across the *B. cinerea* isolates, and that this collection of *B. cinerea* isolates from tomato do not display a strong host-specificity for tomato (Martinez, Blancard et al. 2003, Ma and Michailides 2005, Rowe and Kliebenstein 2007, Samuel, Veloukas et al. 2012).

### Pathogen Specialization to Host Variation

Though we did not find evidence for *B. cinerea* adaptation to tomato based on isolate host source, the *B. cinerea* isolates may contain genetic variation at individual loci that allow them to better attack subsets of the tomato genotypes (Rowe and Kliebenstein 2007, Kretschmer and Hahn 2008, Corwin, Subedy et al. 2016). A visual analysis of the data suggested an interaction between the genomes of *B. cinerea* and tomato (Figure 1 c-h). However, when using the full model, we found no significant interaction between isolate and individual host genotype, even though there was a large fraction of variance within these terms (Table 1). This may indicate a lack of interaction between genetic variation in the host and pathogen. However, this negative result may also be because F-tests in factors with high degrees of freedom can be underpowered, as in the case of the isolate by plant genotype interaction term with 940 degrees of freedom (Table 1). To assess these possibilities, we used an additional statistical approach to test for an interaction between *B. cinerea* and host genotype. We used a Wilcoxon signed-rank test to test if the rank of *B. cinerea* isolate-induced lesion size significantly changes between pairs of tomato genotypes. This showed that when using the full isolate population, the rank performance of the isolates does significantly vary between host genotypes. When comparing mean lesion size between paired plant genotypes, 58% (38 out of 66) of tomato accession pairs had significantly different ranking of the isolates (Wilcoxon signed-rank test with FDR-correction, Table 2, Figure S1, Table S2). A significant p-value indicates that the two host genotypes show evidence for different virulence interactions with the population of *B*. *cinerea* isolates, providing evidence for host x pathogen genotypic interactions. This pattern was consistent across domesticated host pairs, wild host pairs, or between-species host pairs (Wilcoxon signed-rank test with FDR-correction, Table 2). This suggests that the population of *B. cinerea* does display differential responses to the tomato genetic variation.

To focus on whether specific *B. cinerea* isolates may be sensitive to domestication, we applied a Wilcoxon and ANOVA approach. Overall, most isolates (78/97, 80%) are more virulent on domesticated than wild tomato (Figure 3). The Wilcoxon signed-rank test, to compare the rank of mean lesion size of all the *B. cinerea* isolates on wild versus domesticated tomato, was significant (Wilcoxon signed-rank test, W = 5946, p-value = 0.002) (Figure 3). To identify the pathogen genotypes most sensitive to domestication, we conducted single-isolate ANOVAs including the fixed effects of plant, domestication, and experiment, and found two isolates with a significant effect of domestication on lesion size (p < 0.05, FDR corrected) (Figure 1h), both of which are more virulent on domesticated tomato. These included one of the highly virulent isolates (Fd2), and one of the largely saprophytic isolates (Rose), which suggests that isolate virulence level on tomato does not predict *B. cinerea* genetic response to tomato domestication. Both of these isolates were more virulent on domesticated than on wild tomato. These results suggest that this *B. cinerea* population contains two highly domestication-sensitive isolates which are more virulent on domesticated tomato, and a broader pattern of *B. cinerea* specialization to tomato domestication.

### Quantitative Genetics of Pathogen Virulence on Tomato

Genetic variation within *B. cinerea* had a large effect on virulence on tomato and interacted with tomato domestication (Table 1). This suggests that there is genetic variation within the pathogen in which some alleles enhance and other alleles decrease virulence depending upon the plant’s genotype. To identify variable pathogen genes controlling differential virulence across plant genotypes, we conducted a GWA mapping analysis within the pathogen. Due to the large effect of plant genotype on resistance to *B. cinerea*, we performed GWA using the model-corrected least-squared mean virulence measured on each tomato genotype as separate traits. We used a ridge-regression approach in combination with 272,672 SNPs from *B. cinerea* to estimate the phenotypic effects across the genome (Shen, Alam et al. 2013, Corwin, Copeland et al. 2016, Corwin, Subedy et al. 2016, Francisco, Joseph et al. 2016). To determine significance of SNP effects, we permuted phenotypes 1000 times to calculate 95, 99, and 99.9% effect size thresholds within each plant host. This GWA analysis showed that the genetic basis of *B. cinerea* virulence on tomato is highly polygenic. We identified from 1,284 to 25,421 SNPs within *B. cinerea* that were significantly associated with altered virulence on the 12 different host genotypes (significance was determined by the SNP effect size estimate exceeding the 99% permutation threshold using 10,000 permutations). There were no SNPs with large effect sizes, showing the polygenic nature of the trait in the pathogen (Figure 4).

**Figure 4.**
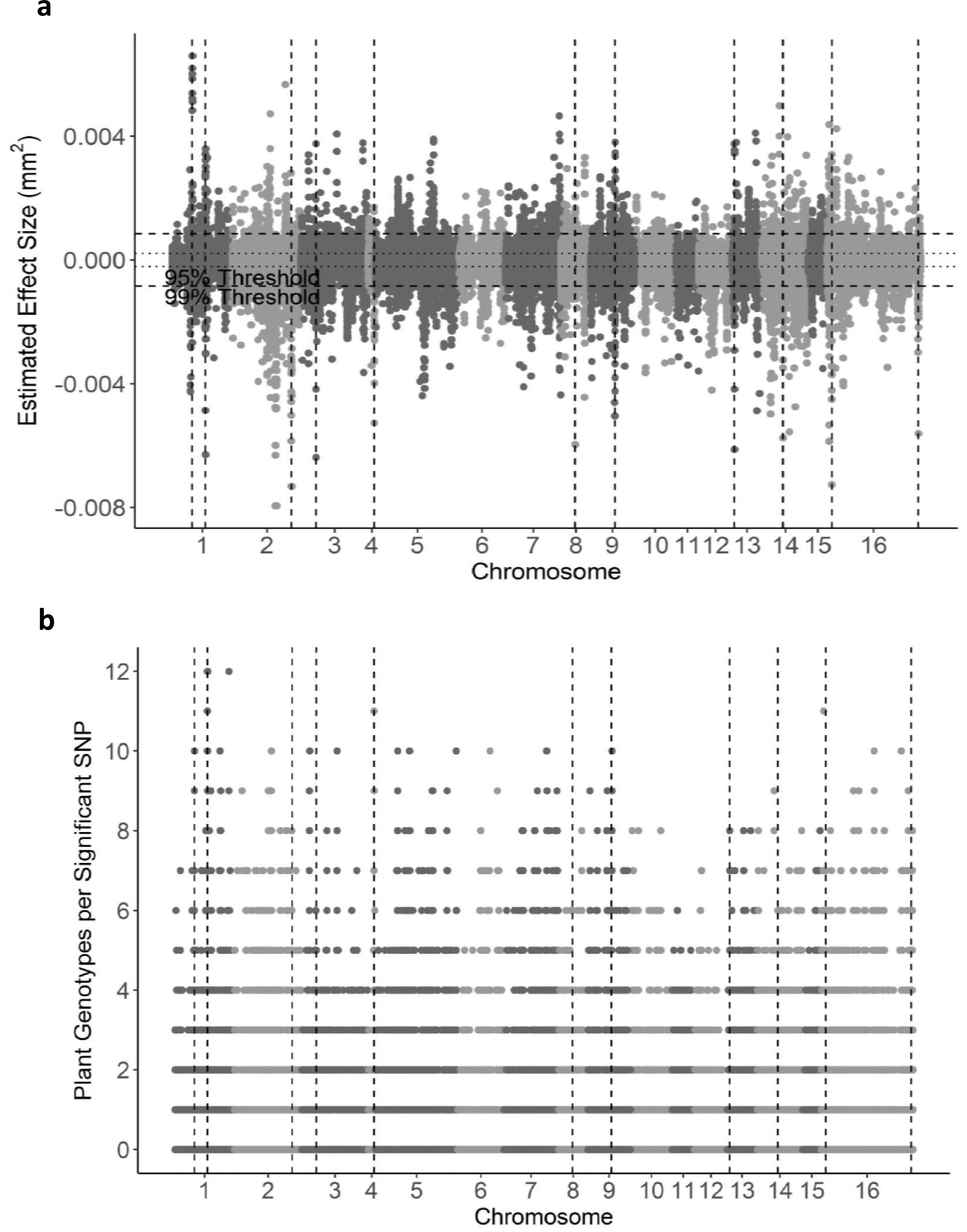
GWA of *B. cinerea* lesion size on individual tomato genotypes. *Botrytis cinerea* chromosomes are differentiated by shading, alternating black and grey. a) Manhattan plot of estimated SNP effect sizes for *B. cinerea* lesion size using a single tomato accession, LA2093. Permutation-derived thresholds are shown in horizontal dashed lines. b) The number of tomato accessions for which a *B. cinerea* SNP was significantly linked to lesion development using the 99% permutation threshold. Frequency is number of phenotypes in which the SNP exceeds the threshold. Vertical dotted lines identify regions with overlap between the top 100 large-effect SNPs for LA2093 and significance across the majority (≥6) of tomato genotypes tested.

While only a small subset of these *B. cinerea* SNPs were linked to virulence on all the tomato genotypes, we were able to obtain better overlap by focusing on gene windows. We found five *B. cinerea* SNPs significantly linked to altered lesion size on all 12 tomato accessions (Figure 4b). 215 SNPs were called in at least ten hosts, and 3.3k SNPs were called in at least half of the hosts while 27% (46,000) of the significant SNPs were linked to virulence on only a single host tomato genotype. These levels of overlap exceed the expected overlap due to random chance (Figure 5a). To change from a SNP-by-SNP focus to a gene-centric focus, we classified a gene as significantly associated if there was 1 SNP linked to a trait using a 2kbp window surrounding the start and stop codon for a given gene. This analysis identified 6 genes linked to differential virulence in all 12 tomato accessions (Figure 5b, Table S3), as some SNPs within a gene had accession-specific phenotypes (significant in <12 tomato accessions). A further 233 genes were linked to differential virulence on between 7 and 11 tomato accessions (Figure 5b, Table S3). Of the 6 genes with SNPs significantly associated with *B. cinerea* virulence on all tomato genotypes, two are heterokaryon incompatibility loci (Bcin01g10020; BcT4_2485), one is a major facilitator superfamily gene, and the remaining 3 are enzymes (peptidase dimerization, Bcin01g10130; pectinesterase, Bcin14g00870; protein kinase, Bcin15g04110). Four of those genes also represent significantly overrepresented functional annotation categories; including heterokaryon incompatibility, pectinesterase, peptidase dimerization, and protein kinase. While most of these genes have not been formally linked to pathogen virulence, pectinesterases are key enzymes for attacking the host cell wall, suggesting that variation in this pectinesterase locus and the other loci may influence pathogen virulence across all the tomato genotypes (Valette-Collet, Cimerman et al. 2003).The SNPs within the pectinesterase gene (BcT4_6001, Bcin14g00870) were only associated with at most 11 tomato accessions while the gene itself is associated with altered virulence on all tomato accessions. This suggested that there may be multiple haplotypes in this locus linked to virulence. To visualize the SNP effects across a single gene and look for evidence of multiple haplotypes, we plotted the effect sizes for all SNPs in this gene and investigated the linkage disequilibrium amongst these SNPs (Figure 6). This showed that the effect of SNPs across this gene vary in effect direction depending on tomato host genotype (Figure 6a), and that there appear to be two different haplotype blocks contributing to the association of this gene to the virulence phenotype (Figure 6b). One block is associated with SNPs in the 5’ untranslated region in SNPs 5-11 and the second block is SNPs that span the entirety of the gene in SNPs 13-26. Interestingly, there are only two SNPs in the open reading frame of the associated gene (Figure 6). This suggests that the major variation surrounding this locus is controlling the regulatory motifs for this pectinesterase. Thus, there is significant genetic variation in *B. cinerea* virulence that is dependent upon the host’s genetic background. This suggests that the pathogen relies on polygenic small effect loci, potentially allowing selection to customize virulence on the different tomato hosts.

**Figure 5.**
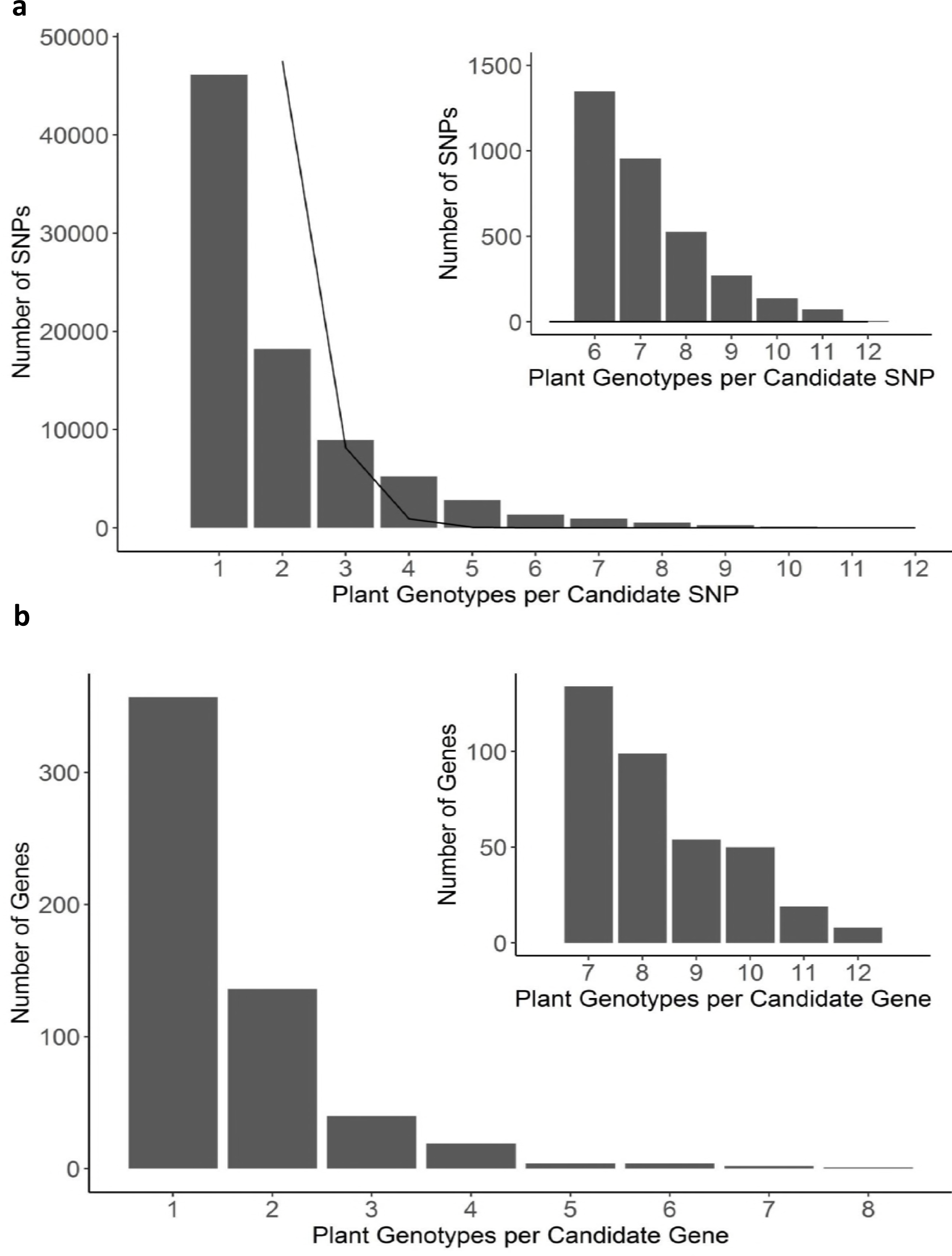
Frequency of overlap in *B. cinerea* GWA significance across tomato accessions. a) Frequency with which the *B. cinerea* SNPs significantly associated with lesion size on the 12 tomato accessions using the 99% permutation threshold. Black lines indicate the expected frequency of overlap, given the number of significant SNPs per plant genotype and size of total SNP set. b) Frequency with which a *B. cinerea* gene significantly associated with lesion size on the 12 tomato accessions. Genes were called as significant if there was one significant SNP in the top 1000 called at the 99% permutation threshold within the gene body, or within 2kb of the gene body.

**Figure 6.**
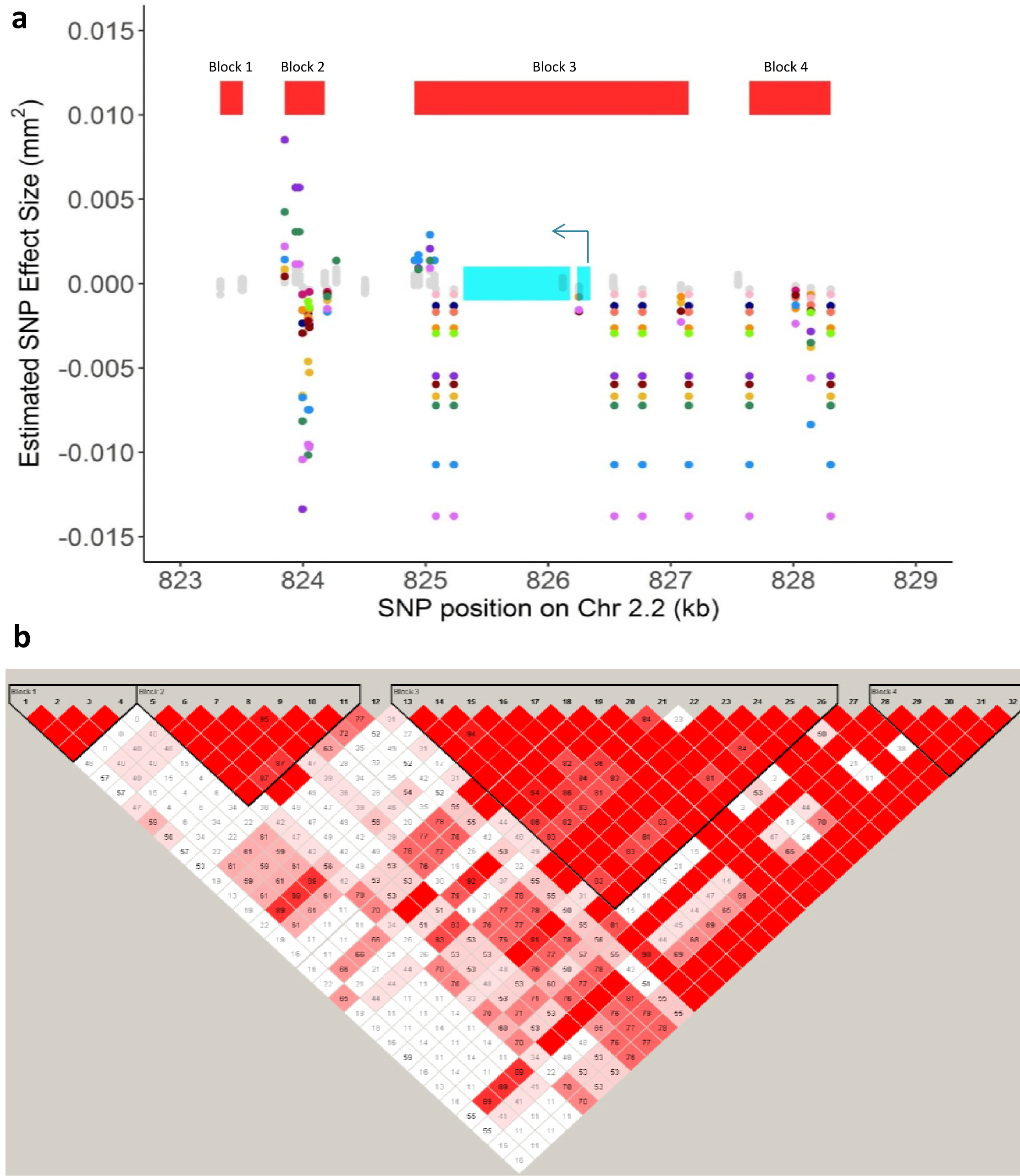
Host specificity of significant SNPs linked to the gene BcT4_6001 (Bcin14g00870). a) SNPs with effects estimates above the 99% permutation threshold are colored by trait (plant phenotype in which the effect was estimated). BcT4_6001 (Bcin14g00870) is a pectinesterase gene linked to at least one significant SNP on all 12 of the tested tomato accessions. The annotated exons are depicted as turquoise rectangles, with the start codon marked with an arrow indicating the direction of transcription. Red rectangles indicate corresponding linkage disequilibrium blocks from Figure 6b. b) Linkage disequilibrium plot, including all pairwise comparisons of SNPs in the 2kb region surrounding Bcin14g00870. The color scheme for each SNP pair is D'/LOD: white if LOD <2 and D’ <1, bright red for LOD ≥2 and D’=1, intermediate shades for LOD≥2 and D’<1.

### Quantitative Genetics of Pathogen Response to Tomato Domestication

The identification of two isolates that distinctly respond to tomato domestication suggests that there is natural genetic variation in *B. cinerea* that is affected by tomato domestication. To directly map *B. cinerea* genes that control differential virulence on wild versus domestic tomatoes, we used the least-squared mean virulence of each isolate across all wild and all domesticated tomato genotypes as two traits. We also calculated a domestication sensitivity trait; the relative difference in lesion size for each isolate between domesticated and wild hosts. Using these three traits, we conducted GWA within *B. cinerea* to map genes in the pathogen that respond to domestication shifts in the plant. Using the mean lesion area of the *B. cinerea* isolates on the wild or domestic tomato hosts identified a complex pattern of significant SNPs similar to the individual tomato accessions (Figure 4, Figure 7). This had a high degree of overlap between the two traits. In contrast, the Domestication Sensitivity trait identified a much more limited set of SNPs that had less overlap with either the mean lesion area on Domesticated or Wild tomato (Figure 7). To begin querying the underlying gene functions for these various *B. cinerea* loci, we called genes as significant if there was one SNP within 2kb of that gene (Figure 7c). Using all 1251 genes linked to domestication phenotypes for a functional enrichment analysis found only 22 significantly overrepresented biological functions (Fisher exact test, p<0.05, Table S3) when compared to the whole-genome annotation. Of the 22 functions overrepresented for domestication virulence traits, eight are enzymes and two are transporters (Table S3). Eight gene functions are uniquely overrepresented in *B. cinerea* growth on wild tomato genotypes, and eight functions are overrepresented only for domestication-sensitivity genes. Among the eight gene functions associated specifically to domestication-sensitivity is indoleamine 2,3-dioxygenase, which converts tryptophan to N-formylkyneureine and has been linked to altered immune responses in a number of systems (Uyttenhove, Pilotte et al. 2003, Chen, Liang et al. 2008, Camañes, Scalschi et al. 2015). The only other known function is a phosphodiesterase related to BcPde2, a gene that has previously been associated with *B. cinerea* virulence through the cAMP signaling pathway (Harren, Brandhoff et al. 2013). Thus, there is an apparent subset of *B. cinerea* genes that may be specific to the genetic changes that occurred in tomato during domestication. Further work is needed to assess if and how variation in these genes may link to altered virulence on domestic and wild tomatoes.

**Figure 7.**
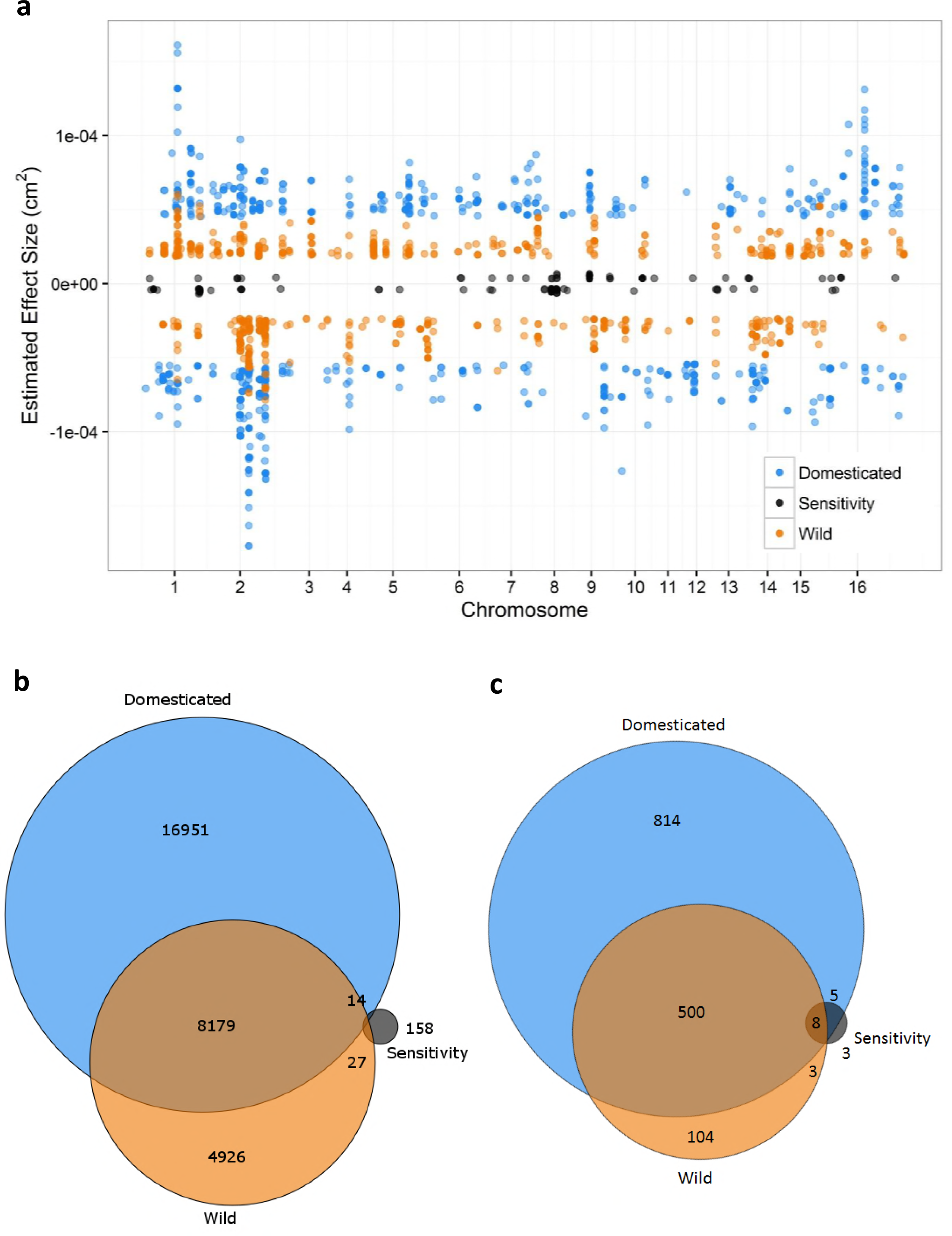
GWA analysis of domestication sensitivity in *B. cinerea*. Domestication sensitivity of each isolate was estimated using the average virulence on the wild and domesticated tomato germplasm and using calculated Sensitivity. This was then utilized for GWA mapping. a) The top 1000 SNPs that significantly affect lesion size across domesticated tomato, wild tomato or domestication sensitivity are shown. Significance is called as crossing the 99% permutation threshold. b) Venn diagram of overlapping SNPs identified as crossing the 99% permutation threshold for each trait. c) Venn diagram of overlapping genes identified as crossing the 99% permutation threshold for each trait. Genes were called as significant if there was one significant SNP within the gene body or within 2kb of the gene body.

## DISCUSSION

The genetics of plant resistance to generalist pathogens are mostly quantitative, depend upon pathogen isolate, and rely on genetic variation in both signal perception and direct defense genes (Kover and Schaal 2002, Parlevliet 2002, Glazebrook 2005, Nomura, Melotto et al. 2005, Goss and Bergelson 2006, Tiffin and Moeller 2006, Rowe and Kliebenstein 2008, Barrett, Kniskern et al. 2009, Corwin, Copeland et al. 2016). Previous studies on tomato resistance to *B. cinerea* have found a quantitative genetic architecture that varies between domesticated and wild tomato species, with higher resistance in the wild species (Egashira, Kuwashima et al. 2000, Nicot, Moretti et al. 2002, Guimaraes, Chetelat et al. 2004, Finkers, van Heusden et al. 2007, Ten Have, van Berloo et al. 2007, Finkers, Bai et al. 2008). However, it was not known how the choice of *B. cinerea* isolate may change this plant-pathogen interaction. To address these questions, we used genetic variation in wild and domesticated tomato accessions in conjunction with a population of *B. cinerea* isolates. This also allowed us to test how domestication within tomato influenced the interaction at the level of the pathogen population and individual genes in the pathogen. *B. cinerea* virulence on tomato, as measured by lesion size, was significantly affected by pathogen isolate, host genotype, and domestication status (Table 1). Tomato domestication led to a slight but significant decrease in resistance to the pathogen but critically, there was no evidence of a domestication bottleneck, with the wild and domesticated tomato accessions having similar variance in resistance (Table 1, Figure 2). There was also little evidence in this *B. cinerea* population for specialization to tomato, supporting the hypothesis that *B. cinerea* is a generalist at the isolate and species level (Figure 1 c-h) (Giraud, Fortini et al. 1999, Martinez, Blancard et al. 2003, Ma and Michailides 2005). GWA mapping within the pathogen showed that the genetics underlying *B. cinerea* virulence on tomato are highly quantitative, and vary across tomato genotypes and domestication status (Figure 5, Figure 7). This analysis identified a small subset of pathogen genes whose variation contributes to differential virulence on most of the hosts tested, and a set of pathogen genes whose variation is responsive to tomato domestication (Table S3).

### Domestication and altered pathogen virulence genetics

These results provide evidence of a mild tomato domestication effect on resistance to the generalist pathogen, *B. cinerea.* We measured an 18% increase in susceptibility across domesticated varieties, but this represents less than 1% of the total variance of *B. cinerea* lesion size on tomato (Table 1). As such, domestication status alone is a poor predictor of a specific tomato host’s resistance to infection by *B. cinerea*. This suggests that while tomato domestication does affect this plant-pathogen interaction, it is not the primary factor defining the measured trait. The effect of tomato domestication varied across the *B. cinerea* isolates, with specific isolates and loci linked to differential virulence across wild and domestic tomatoes (Figure 1 c-h). If a study relies on one or a few isolates, it could obtain a falsely high or falsely low estimation of how host domestication influences pathogen resistance. This shows the need to utilize a population of *B. cinerea* to understand the factors contributing to *B. cinerea* virulence and how this is altered by crop domestication.

In biotrophic pathogens, host domestication has decreased the diversity of resistance alleles because they are lost in the domestication bottleneck as found for specialist pathogens (Tanksley and McCouch 1997, Doebley, Gaut et al. 2006, Hyten, Song et al. 2006, Chaudhary 2013). Surprisingly, we did not find evidence for a domestication bottleneck in the phenotypic resistance to *B. cinerea* (Figure 2, Figure 3). This is in contrast to genomic studies that explicitly show a genotypic bottleneck within tomato domestication (Miller and Tanksley 1990, Koenig, Jiménez-Gómez et al. 2013). This suggests that at least for this generalist pathogen, the genetic bottleneck of tomato domestication has not imparted a phenotypic bottleneck. One possible explanation is that resistance to this pathogen is so polygenic in the plant that our experiment is not sufficiently large to pick up any genetic bottleneck effect using phenotypic variance. These patterns, of mild decrease in resistance to *B. cinerea* due to plant domestication, and within-species plant variation exceeding the contribution of domestication itself, may be unique to interactions between *B. cinerea* and tomato, or more general. It remains to be seen if these patterns hold for *B. cinerea* on its other host plants. It is unclear whether domestication has a universal effect on plant resistance to *B. cinerea*, or if each domestication event is unique.

### Polygenic quantitative virulence and breeding complications

Our results indicate a highly polygenic basis of quantitative virulence of the generalist *B. cinerea* on tomato. The variation in lesion size is linked to numerous *B. cinerea* SNPs, each with small effect sizes (Figure 4a). Importantly, the tomato host accession greatly influenced which *B. cinerea* loci were significantly associated to lesion size (Figure 5). Thus, it possible that there is specialization at the gene level, in which different alleles within the pathogen link to differential virulence on specific host genotypes (Giraud, Fortini et al. 1999, Rowe and Kliebenstein 2007, Blanco-Ulate, Morales-Cruz et al. 2014). This polygenic architecture of virulence is distinctly different from specialist pathogens that often have one or a few large effect genes that control virulence (Keen 1992, De Feyter, Yang et al. 1993, Abramovitch and Martin 2004, Boyd, Ridout et al. 2013, Vleeshouwers and Oliver 2014). Further studies are needed to compare how the host plant species may affect this image of genetic variation in virulence.

These results indicate some particular challenges for breeding durable resistance to *B. cinerea* and possibly other generalist pathogens. The highly polygenic variation in virulence combined with genomic sequencing, showing that this pathogen is an inter-breeding population, suggests that the pathogen is actively blending a large collection of polymorphic virulence loci (Rowe and Kliebenstein 2007, Fekete, Fekete et al. 2012, Atwell, Corwin et al. 2015, Atwell, Soltis et al. 2017). Thus, it is not sufficient to breed crop resistance against a single isolate of *B. cinerea*, as this resistance mechanism would likely be rapidly overcome by new genotypes within the field population of *B. cinerea*. In contrast, it is likely necessary to breed resistance using a population of the pathogen, and to focus on plant loci that target entire virulence pathways or mechanisms. The results in this study indicate that the specific genetics of the plant host, the general domestication status, and the specific genetics of the pathogen isolate will all combine to affect how the estimated breeding value inferred from any experiment will translate to a field application (Table 1). As such, utilizing a single or even a few pathogen isolates to guide resistance breeding in plants is unlikely to translate to durable resistance against *B. cinerea* as a species. However, the lack of a domestication bottleneck on tomato resistance to B*. cinerea* suggests that, at least for tomato, allelic variation in this generalist pathogen is sufficient to overcome introgression of wild resistance genes or alleles into the domesticated crop.

### Conclusion

This study examined the contributions of host and pathogen natural genetic variation to the quantitative interaction in the tomato-*B. cinerea* pathosystem. In addition, the study explicitly tested the effects of tomato domestication on this pathosystem. *B. cinerea* has a highly quantitative genetic basis of virulence on tomato, which is dominated by pathogen effects but also sensitive to genetic variation linked to tomato domestication. Future studies are necessary to test if this pattern of domestication responses in tomato is similar to what happens in other crops. Because this population of *B. cinerea* can infect a wide range of hosts, it will be possible to directly conduct this study. By extending future work to additional domestication events, it may be possible to test if independent crop domestication events have a consistent underlying genetic signal of *B. cinerea* adaptation to plant domestication.

